# Active membrane conductances and morphology of a collision detection neuron broaden its impedance profile and improve discrimination of input synchrony

**DOI:** 10.1101/454702

**Authors:** Richard B. Dewell, Fabrizio Gabbiani

## Abstract

**Abstract:** Brains processes information through the coordinated efforts of billions of individual neurons, each encoding a small part of the overall information stream. Central to this is how neurons integrate and transform complex patterns of synaptic inputs. The neuronal membrane impedance sets the gain and timing for synaptic integration, determining a neuron’s ability to discriminate between synaptic input patterns. Using single and dual dendritic recordings *in vivo*, pharmacology, and computational modeling, we characterized the membrane impedance of a collision detection neuron in the grasshopper, *Schistocerca americana*. We examined how the cellular properties of the lobula giant movement detector (LGMD) neuron are tuned to enable the discrimination of synaptic input patterns key to its role in collision detection. We found that two common active conductances g_H_ and g_M_, mediated respectively by hyperpolarization-activated cyclic nucleotide gated (HCN) channels and by muscarine sensitive M-type K+ channels, promote broadband integration with high temporal precision over the LGMD’s natural range of membrane potentials and synaptic input frequencies. Additionally, we found that the LGMD’s branching morphology increased the gain and decreased delays associated with the mapping of synaptic input currents to membrane potential. We investigated whether other branching dendritic morphologies fulfill a similar function and found this to be true for a wide range of morphologies, including those of neocortical pyramidal neurons and cerebellar Purkinje cells. These findings further our understanding of the integration properties of individual neurons by showing the unexpected role played by two widespread active conductances and by dendritic morphology in shaping synaptic integration.

*New & Noteworthy:* Neuronal integration depends on complex interactions between synaptic input patterns and the electrochemical properties of dendrites. We used an identified collision detection neuron in grasshoppers to examine how its morphology and two membrane conductances determine the gain and timing of synaptic integration in relation to the computations it performs. The neuronal properties examined are ubiquitous and therefore further a general understanding of neuronal computations, including those in our own brain.

## Introduction

By the nineteen forties, neuroscientists had recognized that neuronal membranes exhibited a frequency dependent impedance and had begun investigating which cellular mechanisms contributed to their resistance, capacitance, and inductance (Cole 1941). The capacitance of the membrane is determined by its lipid bilayer, and its resistance by permeability to ion fluxes, but the inductance came from an unknown source. After the discovery of active conductances, it became clear that their kinetics produced a phenomenological inductance, and that its properties could be investigated using Hodgkin and Huxley’s equations (Mauro 1961; Mauro et al. 1970).

Membrane capacitance low-passes, while inductance high-passes input currents. Their combination can thus result in bandpass filtering and resonance. In the decades since, researchers have learned a great deal about the roles of active conductances in shaping the electrical properties of neurons, and the influence of neuronal bandpass properties on rhythmic activity within neural networks (Das et al. 2017; Hutcheon and Yarom 2000; Wang 2010). In addition to generating neural rhythms, the membrane impedance shapes the integration properties of dendrites determining their ability to discriminate between patterns of synaptic inputs.

Voltage-gated channels modify the membrane impedance through their time varying conductances. The range of input frequencies affected depends on the channel kinetics. Non-inactivating channels that decrease their conductance as the membrane potential approaches their reversal potential produce negative feedback on changes in membrane potential and thus a phenomenological inductance. In the LGMD two such inductive channels, HCN and M, produce the largest resting conductances, influencing the selectivity and timing of its responses (Dewell and Gabbiani 2018a, 2018b). The impedance profile of a neuron is also influenced by its dendritic morphology. An extended morphology causes dendritic compartmentalization which increases the attenuation and lag of postsynaptic potentials traveling towards the spike initiation zone, but simultaneously enhances local coincidence detection and nonlinear processing (Häusser and Mel 2003; London and Hausser 2005). Dendritic branching and tapering could optimize synaptic integration by increasing the spatial homogeneity of postsynaptic potentials’ influence at the site of spike initiation (Cuntz et al. 2007). Additionally, the dendritic morphology of pyramidal and Purkinje neurons is believed to favor the neurons’ response to inputs of certain frequencies thus promoting neuronal resonance (Dhupia et al. 2015; Ostojic et al. 2015).

We have chosen to explore these questions in the LGMD (O’Shea and Williams 1974), an identified neuron in locusts that selectively responds to visual stimuli mimicking impending collision (Rind and Simmons 1992; Schlotterer 1977). The LGMD’s stimulus selectivity arises from the precise spatiotemporal patterning of synaptic inputs (Jones and Gabbiani 2010; Peron et al. 2007a; Zhu and Gabbiani 2016) and complex dendritic computations, including filtering by active conductances such as sodium, low voltage calcium, calcium-dependent potassium, inactivating potassium, HCN, and M channels (Dewell and Gabbiani 2018a, 2018b; Gabbiani et al. 2002; Peron and Gabbiani 2009). Different aspects of the LGMD’s firing patterns – including burst firing – have been tied to the generation of escape behaviors (Dewell and Gabbiani 2018a; Fotowat et al. 2011; McMillan and Gray 2015). Currently, no one has characterized the LGMD’s membrane impedance to determine how it shapes this neuron’s dendritic integration and visual computations.

Neuronal membrane properties have mainly been studied with cultured neurons or brain slices and the natural synaptic input patterns are still unknown for most neurons, so much remains unanswered about how the impedance properties of neurons influence synaptic integration and neural computations *in vivo*. As both g_H_ and g_M_ are sensitive to numerous modulators (Delmas and Brown 2005; Wahl-Schott and Biel 2009), our ability to conduct experiments *in vivo* ensures that the channels are in the relevant modulatory state. We discovered that the LGMD membrane impedance is broadband and generates small delays between synaptic current and membrane potential that remain consistent across membrane potentials. The HCN and M channels, their interactions, and the neuronal morphology all contribute to producing this broadband impedance profile which increases the LGMD’s ability to discriminate behaviorally relevant patterns of synaptic inputs from irrelevant ones.

## Materials and Methods

### Animals

Experiments were performed on adult female grasshoppers 7-12 weeks of age (*Schistocerca americana*). Animals were reared in a crowded laboratory colony under 12 h light/dark conditions. Animals were selected for health and size without randomization, and investigators were not blinded to experimental conditions. Sample sizes were not predetermined before experiments. The surgical procedures used have been described previously (Dewell and Gabbiani 2018a; Gabbiani and Krapp 2006a; Jones and Gabbiani 2012).

### Visual stimuli

Visual stimuli were generated using Matlab (RRID:SCR_001622) and the Psychophysics Toolbox (PTB-3, RRID:SCR_002881) on a personal computer running Windows XP. A conventional cathode ray tube monitor refreshed at 200 frames per second was used for stimulus display (LG Electronics, Seoul, Korea). Looming stimuli are the two-dimensional projections of an object approaching on a collision course with the animal. They consisted of an expanding dark square simulating a solid object with half size *l* approaching at constant speed *ν* (illustrated in Figure 1B). The expansion profile is characterized by the ratio *l*/|*ν*|, as previously described (Gabbiani et al. 2001).

**Figure 1.**
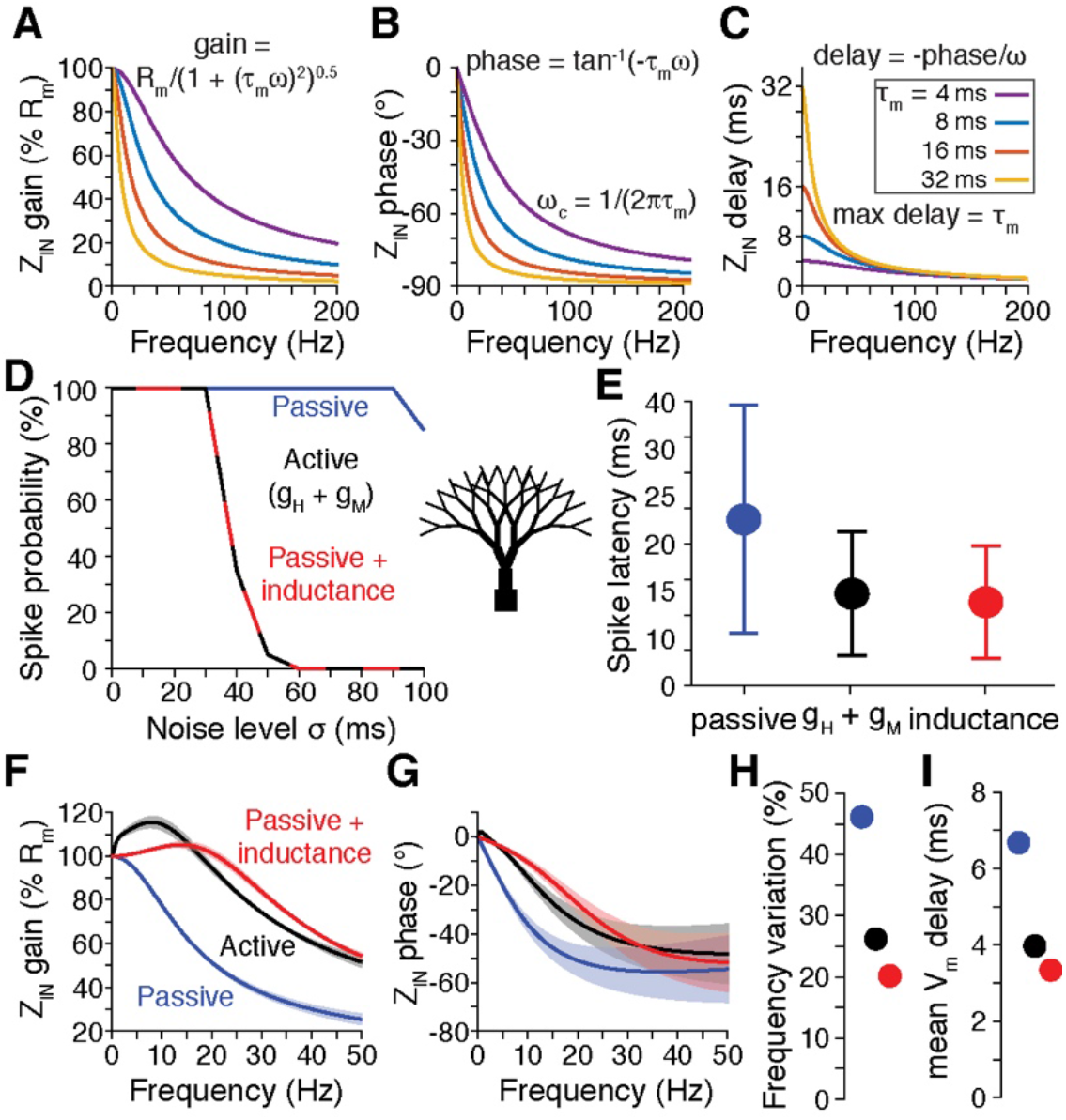
Membrane impedance shapes the ability of neurons to discriminate and transmit the timing of their inputs. A) The membrane impedance amplitude of a passive isopotential neuron (modeled as an RC circuit) decays with frequency, more rapidly so the slower the membrane time constant (τ_m_). The impedance amplitude determines the gain between input currents and membrane potential. Formula: R_m_ - membrane resistance; ω-frequency (radian). B) The impedance phase, which determines the timing between input currents and membrane potential, also decreases with frequency in passive isopotential neurons. The corner frequency (ω_c_) of the membrane impedance for such RC models is determined by the τ_m_. C) Converting the phase into the time domain reveals that for low frequency signals the delay between membrane current and potential equals τ_m_ and that this delay depends on frequency. D) Simulations with a Rall branching neuron model (schematic at right) tested the ability to discriminate the synchrony of 100 synaptic inputs. If the model contained the active conductances g_H_ and g_M_, spikes were generated by synchronous inputs but not if input synchrony was decreased by adding random noise to the synaptic activation times (black). Removal of the active conductances greatly decreased the ability to discriminate the timing of synaptic inputs (blue). If the voltage dependent conductances were replaced by a voltage independent inductance, the ability to discriminate input timing was returned. E) The timing of spikes changed accordingly, such that for the models with inductances the spike time more faithfully conveyed the timing of its inputs. F,G) The voltage independent inductance produced similar changes to the impedance gain (F) and impedance timing (G) as did the voltage dependent conductances. H) The passive model had higher frequency variation indicating that it was more lowpass and less broadband. I) The mean absolute delay between I_m_ and V_m_ was higher for the passive model, causing the reduction in timing reliability for the model.

### Electrophysiology

Sharp-electrode LGMD intracellular recordings were carried out in current-clamp or voltage-clamp mode using thin walled borosilicate glass pipettes (outer/inner diameter: 1.2/0.9 mm; WPI, Sarasota, FL; see Jones and Gabbiani, 2012, as well as Dewell and Gabbiani, 2018a for details). The membrane potential (V_m_) and current (I_m_) were low-pass filtered with cutoff frequencies of 10 kHz and 5 kHz, respectively. Most recordings were digitized at a sampling rate of 20,073 Hz (for some experiments V_m_ was digitized at 40,146 Hz). We used a single electrode clamp amplifier capable of operating in discontinuous mode at high switching frequencies (20-35 kHz; SEC-10, NPI, Tamm, Germany). Responses to current injections were recorded in discontinuous current clamp mode (DCC). For dual recordings we inserted under visual guidance a second sharp electrode into the excitatory dendritic field of the LGMD (see Figure 3A) with a motorized micromanipulator (Sutter Instruments, Novato, CA). Membrane potential was recorded with a second SEC-10 amplifier in bridge mode with electrode resistance and capacitance compensation when not injecting current and in DCC mode while injecting current. Switching frequencies, signal filtering and digitization were identical for both recordings.

The physical distance along the dendrites and an estimate of the electrotonic distance between recording electrode pairs were measured as follows. First, all cells were stained with Alexa Fluor 594 and imaged with a CCD camera to record electrode positions (Dewell and Gabbiani 2018a). Next, an image of the neuron and electrodes (cf. Figure 3A) was imported into ImageJ (RRID:SCR_003070), the dendritic path between the electrode tips was manually traced, and the corresponding path length recorded. To estimate the electrotonic distance we measured the amplitude of the back-propagating action potentials (bAP) at each location and used the difference in these amplitudes. As in a previous study, the differences in bAP amplitude provided a more reliable explanatory variable for distance dependent effects (Dewell and Gabbiani 2018a).

A neuron’s frequency-dependent membrane properties are characterized by its impedance profile. The subthreshold impedance can be decomposed into input impedance which describes local change in membrane potential at the site of an input current and transfer impedance that describes change in membrane potential at remote locations after propagation through the neuron. Both input and transfer impedance shape synaptic integration within dendrites. Additionally, impedance properties at the site of spike initiation influence the transformation of membrane potential into spiking activity, including the frequency filtering and timing of a neuron’s output. If a neuron’s conductance distribution is uniform, then the transfer and input impedances will have similar properties. When it is not, the impedance profile varies by location and transfer depends on the recording locations and the direction of signal propagation (Hu et al. 2009; Vaidya and Johnston 2013).

To measure the impedance profile, we injected sine waves of increasing frequency called chirp (or zap) currents. We recorded the injected membrane current simultaneously with the membrane potential to avoid any discrepancies between the computer-generated waveform and the current injected by the amplifier. The chirp currents used were identical to those described in Dewell and Gabbiani (2018b). The chirp current is defined as *I*(*t*) = *I_p_sinϕ*(*t*), where *I_p_* is the peak current, *ϕ*(*t*) is the phase of the sine wave, and its instantaneous frequency is defined as 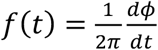 (in units of Hz). We generally used chirps with a duration of 20 s (sometimes 30 s, as is illustrated in Figure 3B) that increased in frequency either linearly or exponentially with time. In most experiments, as well as for all simulations, we used exponentially increasing chirp currents. The linear chirp started at 0 Hz and was calculated as *I*(*t*) = *I_p_*sin (*πβt*^2^), with *t* being the time from the start of the chirp (in units of s) and *β* the rate of increase in instantaneous chirp frequency (in Hz/s). The exponential chirp was given by *I*(*t*) = *I_p_*sin (2*πf*_0_*te^βt^*), where *f*_0_ is the initial chirp frequency and *β* determines the (accelerating) rate of frequency increase (Barrow and Wu 2009). For all exponential chirps, we used *f*_0_ = 0.05 Hz and β = 0.24 Hz which produced a chirp current increasing to 35 Hz over 20 s. We saw no differences in the calculated impedance profiles with chirps having different frequency profiles, in accordance with previous comparisons of impedance profiles between chirps of increasing and decreasing frequencies (Ulrich 2002; Erchova et al. 2004; Hu et al. 2002; van Brederode and Berger 2008; van Brederode and Berger 2011) or comparison of linearly increasing chirps and sums of sine wave stimuli (Hutcheon et al. 1996).

### Pharmacology

We applied the HCN-channel blocker ZD7288 and the M-channel blocker XE991 either directly in the bath saline or by local puffing as previously described (Dewell and Gabbiani 2018a, 2018b). For local puffing we used a micropipette connected to a pneumatic picopump (PV830, WPI, Saratoga, FL). Drugs were mixed with physiological saline containing fast green (0.5%) to visually monitor the affected region. For both delivery methods, drug concentrations within the lobula were ~200 μM for ZD7288 and ~100-200 μM for XE991. In dual recording experiments, mecamylamine was applied to block excitatory postsynaptic potentials (EPSPs) and was present in both control and drug conditions. Blockade of EPSPs reduced membrane noise as well as noise in the calculated impedance profiles. In paired comparisons, mecamylamine caused small increases in impedance amplitude (~5%) but this difference was smaller than the variability between animals or between dendritic locations. For control data we thus pooled recordings with and without mecamylamine.

### Experimental design and statistical analyses

Data analysis was carried out with custom MATLAB code (The MathWorks, Natick, MA). Linear fits were done by minimization of the sum of squared errors. Before each chirp current a −2 nA hyperpolarizing step current was used for monitoring the input resistance and the membrane time constant. Unless otherwise specified, all statistical tests were made using the Wilcoxon rank sum test (WRS) which does not assume normality or equality of variance. Normality of the data was assessed by using a Lilliefors test for any tests assuming it.

The complex impedance *Z* was calculated as *Z*(*f*) = fft(V_m_)/fft(I_m_) where fft is the (frequency dependent) fast Fourier transform, I_m_ the membrane current, and V_m_ the membrane potential. Both I_m_ and V_m_ were down sampled to 2 kHz before the fft. *Z* was calculated for each trial and then averaged across trials. The complex impedance is composed of its real part, the resistance *R*, and its imaginary part, the reactance *X*: *Z* = *R* + *i X*. A positive reactance, or inductance, indicates that changes in V_m_ precede changes in I_m_. Conversely, a negative reactance, or capacitance, indicates that changes in I_m_ precede changes in V_m_. The impedance amplitude is calculated as the absolute value of 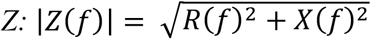. The impedance phase is calculated by the inverse tangent of the reactance divided by the resistance: *Φ*(*f*)= tan^-1^*X*(*f*)/*R*(*f*). This impedance phase is equal to the angle between the x-axis and a line from the origin to a point of the impedance locus plot (cf. Figure 5C).

Input impedance (Z_IN_) was calculated from the V_m_ and I_m_ recorded at the site of current injection. Transfer impedance (Z_TR_) was calculated with the I_m_ from the site of current injection and the V_m_ from the non-injected site. Voltage attenuation was calculated as the relative reduction in impedance amplitude from the current injection site to the non-injected site: *V_att_*(*f*) = (|*Z_IN_*(*f*)| − |*Z_TR_*(*f*)|)/|*Z_IN_*(*f*)|. Similarly, the phase lag was calculated as the input minus the transfer impedance phase. The input and transfer impedance phases were each converted to the time domain by *t_phase_*(*f*) = *Φ*(*f*)/(2*πf*) where *f* is the instantaneous chirp frequency (in Hz) calculated as 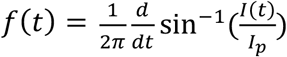, for details see (Dewell and Gabbiani 2018b). The mean V_m_ delay was then calculated by averaging across frequencies the absolute value of the delay between the input current and membrane potential.

Resonant strength was calculated from the input impedance amplitude as the ratio of the maximum impedance to the steady state (0 Hz) impedance. The impedance frequency variation (*f_var_*)was calculated as the standard deviation (across frequencies ranging from 0 to 35 Hz) of the impedance amplitude profile divided by the mean impedance amplitude. Unlike resonance strength, *f_var_* incorporates both the strength and breadth of the bandpass, giving an indication of how ohmic (or broadband) the membrane is. Indeed, an ideal resistor would have a frequency variation of zero while a membrane exhibiting sharp bandpass attributes would produce high frequency variation. The membrane constant values used for our electrical circuit models yielded *f_var_* values of 18.4 for the lowpass circuit, and 378 for the strongest bandpass circuit (see below; Fig. 4G). There is a potential drawback to the statistic, though, since it depends on the frequency range examined, limiting the ability to compare values between studies examining different frequency ranges.

For calculating the power density of the membrane potential, we used the Matlab multitaper power spectral density estimate (function ‘pmtm’), with a time-half bandwidth product of 4 measured over 15 octaves at frequencies of 2^-4^ to 2^11^ Hz with 8 voices (scales) per octave. The mean was subtracted from all data before calculating the power density estimate and the resulting estimates were then averaged across trials. For estimating the time-varying power density we used Matlab’s continuous wavelet transform (function ‘cwt’). The frequency range was set from 0.1 to 2,000 Hz and the signal was not extended. As for the spectral density estimate, wavelet transforms were calculated for individual trials and the resulting wavelet transform coefficients were then averaged across trials. In the calculation of both measures, the raw data was used with no down sampling or filtering.

### Electrical circuit model

The LGMD membrane impedance was compared to the impedance of an equivalent cable based on a series of either RC or RLC circuits. RC circuits describe passive membranes, while RLC circuits can approximate the frequency-dependent properties of active neuronal membranes. Active conductances are both frequency and voltage dependent, while inductances are only frequency dependent. The RLC circuit was modeled as illustrated in Figure 4F. The membrane impedance density was calculated as:

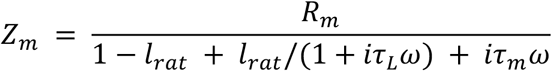

where *Zm* is the input impedance density (in units of Ωcm^2^), *R_m_* is the membrane resistivity (Ωcm^2^), *C_m_* is the membrane capacitance (F/cm^2^), τ_m_ (s) is membrane time constant (τ_m_=*R_m_C_m_*), τ_L_ is the inductance time constant (s), ω is frequency in rad/s, *l_rat_* is the ratio of the membrane conductance that is in series with the inductor (range [0–1]), and *i* = √-1. If *l_rat_* is zero, then no current will pass through the inductor and the circuit is equivalent to an RC circuit with membrane impedance:

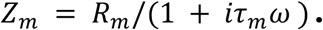

In both cases, the input impedance (Z_IN_) was calculated by dividing *Z_m_* by the unit surface area of the cable (cm^2^). We used the average dendritic circumference of our full LGMD model (20.85 μm) for the cable circumference.

Transfer impedance measures were based upon a cable of fixed diameter and infinite length. The voltage transfer along the cable was calculated as *V_TR_* = *e^−l/λ^*, where *l*(cm) is the distance between the two locations and *λ* is the cable’s length constant (cm), 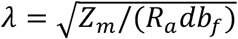. *R_a_* is the axial resistivity (Ωcm), *d* is the diameter (cm), and *b_f_* = 3 is the branch factor — a scalar added to account for additional attenuation due to the dendritic branching. The transfer impedance *Z_TR_* at distance *l* from the site of current injection is the product of *Z_IN_* and *V_TR_*.

### Morphological neuron models

Comparisons between different cell morphologies used published neuron models downloaded from ModelDB (RRID:SCR_007271) and a simplified Rall model (Rall 1964). The Rall model (illustrated in Fig. 1) had 64 sections, each 20 μm long. The soma had a 20 μm diameter and the dendritic arbor had 5 branch levels. The base branch had a 12 μm diameter with each level decreasing in diameter according to the 2/3 power law (Rall 1964). Among the available morphologies, we selected neurons spanning a wide range of branching patterns and sizes. The morphologies used included a cerebellar Purkinje neuron (Miyasho et al. 2001; Ostojic et al. 2015), a CA1 pyramidal neuron (Migliore et al. 2004), an oriens-lacunosum/moleculare (OLM) hippocampal interneuron (Sekulić et al. 2015), and a human layer 2/3 temporal cortex pyramidal neuron (Eyal et al. 2016). As we sought to compare the influence of the cell morphology, all models were passive with identical membrane properties close to those used in the LGMD model (C_m_ = 0.8 μF/cm^2^, G_m_ = 0.1 mS/cm^2^, Ra = 350 Ωcm). All simulations were carried out using the NEURON software simulation package. For a passive model measuring impedance with chirp currents, as was done experimentally, or with NEURON’s built-in impedance tools yielded the same result, so we used the built-in impedance measurement to speed up the simulations carried out for Fig. 8A and B.

In Fig. 8G, the percentage of cell surface area with improved transfer impedance amplitude was calculated from the data depicted in Fig. 8F (right). Specifically, we calculated the transfer impedance amplitude between all neuron segments and compared it to the impedance amplitude of an isopotential model (using the same cell morphology but with axial resistivity set to zero). The impedance amplitude of an isopotential cell is R/(1 +τ_m_^2^ω^2^)^0.5^, where R is the membrane resistance (MΩ), ω is the frequency (s^-1^), and τ_m_ is the membrane time constant (s). As C_m_ and G_m_ were the same for all morphologies, their membrane time constants were the same (τ_m_ = C_m_G_m_) while the membrane resistance changed with their total membrane area A (R = (AG_m_)^-1^). For each segment, we summed the surface area of all segments to which a broadband signal transferred with greater gain than the isopotential value (dashed line in Fig. 8F, right) and divided by the neuron’s total surface area. Frequencies of 0 to 1,000 Hz were used and averaged with equal weight.

Similarly, to obtain Fig. 8I we compared the transfer impedance phase between all segment pairs and compared it to the impedance phase of an isopotential equivalent. The impedance phase of an isopotential cell is tan^-1^(-τ_m_ ω), as illustrated in Fig. 1B, right. For the phase measurements, a weighted average was used with the transfer amplitude between the segment pair determining the weight for each frequency. As signals of some frequencies transfer better than others, the transfer phase of these signals influences the membrane synchrony more. The percentage area with increased impedance phase was then calculated by summing the surface area of segments with more synchronous transfer phase than an isopotential equivalent cell and dividing by the total membrane surface area of the neuron. These calculations effectively measure the average signal transfer of a passive cell with realistic morphology under the simplifying assumptions that broadband inputs impinge upon the neuron with a uniform density. Although these assumptions would not necessarily hold *in vivo* where the locations and frequency of inputs is constantly changing, they provide a method of estimating transfer impedance without relying on the less realistic assumptions that all inputs impinge on a single location or that all signals only integrate at a single location, like the neuron’s spike initiation zone.

## Results

### Broader impedance profiles improve the discrimination of input timing

The input impedance (Z_IN_) of passive neurons can be approximated by RC circuits containing a parallel capacitance and resistance. It is well known that such circuits exhibit a sharp lowpass filtering of input currents. If a sinusoidal current is applied to an RC circuit or a passive neuron, then higher frequencies produce smaller changes in membrane potential resulting in a frequency dependent gain (Fig. 1A). The impedance phase similarly decreases with input current frequency (Fig. 1B). The slower the membrane, or equivalently the higher the membrane time constant, the faster gain and phase decay with frequency. Conversion of the impedance phase to time shows that the delay between an input current and the resulting change in membrane potential is equal to τ_m_ for low frequencies (Fig. 1C). This Z_IN_ delay decreases with increased signal frequency. Long, frequency dependent delays reduce the ability of neurons to discriminate between synaptic input arrival times. To prevent this, auditory neurons that precisely discriminate the timing of their synaptic inputs have small time constants (~0.3 ms), thereby minimizing the delay between input currents and change in V_m_ (McGinley et al. 2012; Mikiel-Hunter et al. 2016; Remme et al. 2014).

Simulations with model CA1 pyramidal neurons show that g_H_ increases the ability of a neuron to discriminate the timing of its synaptic inputs (Migliore et al. 2004). In such simulations, a barrage of synaptic inputs impinges on randomly selected dendritic branches with varying degrees of synchrony. The input synchrony is altered with the addition of a random temporal jitter applied to each input. Replacing g_H_ by an inductance results in a similar increase in synchrony discrimination. To demonstrate this, we simulated 100 synaptic inputs impinging with varying degrees of synchrony onto an artificial model neuron (Fig. 1D; Rall model, Methods). In the passive case, lowpass filtering prevented discrimination of input synchrony (Fig. 1D, blue line). Furthermore, the spike timing was less reliable and had a longer latency (Fig. 1E, blue line). As in the CA1 model simulations, active dendritic conductances, in this case g_H_ and g_M_, increased the discrimination of the input synchrony and produced spikes with shorter, more consistent latencies (Fig. 1D and E, black). Replacing these conductances with an inductance resulted in the exact same increase in input synchrony discrimination with slightly improved spike timing, illustrating the role played by g_H_ and g_M_’s phenomenological inductance (Fig. 1D and E, red).

Among these three models, the passive Rall branching model is the most effective low-pass filter (Fig. 1F,G, blue). The active conductance and the inductance models differ slightly, but share key similarities (Fig. 1F,G). In both models the gain and phase decrease less with frequency than in the passive case, reducing lags at low frequencies. We quantified these two points by computing the frequency variation (Fig. 1H) and the mean V_m_ delay (Fig. 1I). Earlier work showed that discriminating between temporal patterns of excitatory inputs is key to the LGMD’s selectivity for looming stimuli (Jones and Gabbiani 2010). What then is its impedance profile, and can it explain synaptic pattern discriminability in the LGMD?

### The spectral power density of the LGMD’s membrane potential is concentrated at low frequencies

The LGMD possesses three dendritic fields with the largest one, field A, integrating excitatory inputs originating from every facet of the compound eye (Fig. 2A). It responds more or less vigorously to approaching objects that activate thousands of facets based on the object’s trajectory and spatial coherence (Dewell and Gabbiani 2018a; Gray et al. 2001). Currently, it is unknown what synaptic input current frequencies most neurons experience *in vivo*. To address this question, we first measured the power spectrum of the membrane potential for spontaneous activity. The LGMD receives a high number of spontaneous synaptic inputs (Fig. 2D; Jones and Gabbiani, 2012) and its membrane time constant is short *in vivo* (~ 7 ms; Gabbiani and Krapp, 2006; Peron et al., 2007; Dewell and Gabbiani, 2018a, 2018b). This raises the possibility that high frequency components might dominate the LGMD’s membrane potential power spectrum. However, we found that 87% of the power was contained below 35 Hz, with 50% at less than 7.5 Hz (Fig. 2E, black lines).

**Figure 2.**
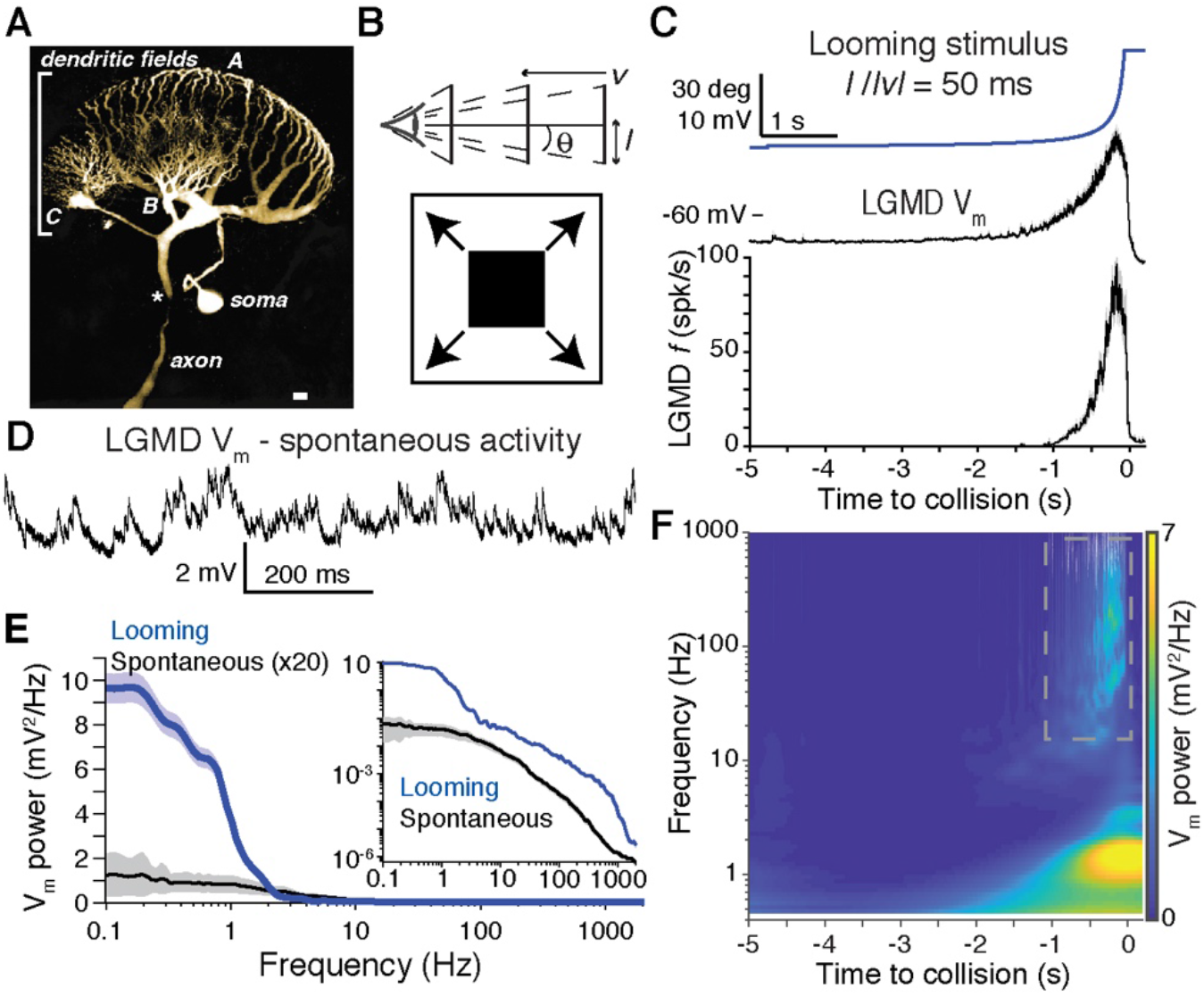
Spontaneous and looming stimulus-evoked spectral power density of the LGMD’s membrane potential. A) Micrograph of the LGMD, illustrating its dendritic fields (labeled A, B and C), the spike initiation zone (SIZ, marked with *) at the start of the axon, and the soma which lies outside of the neuron’s electrical signal path (adapted from Gabbiani et al., 2002). Scale bar: 25 μm. B) A looming stimulus consists of a black square expanding symmetrically on the animal’s retina (bottom). It simulates an object of half size *l* approaching on a collision course at constant velocity *ν* (< 0), subtending an angular size 2*θ*(t) at the retina (top). C) Looming stimuli produce an angular size that increases nonlinearly over time (top, blue line). Most of the response to looming stimuli occurs shortly before the projected time of collision as can be seen in the average membrane potential (V_m_, middle) and firing rate (bottom, displayed as mean ± standard error of the mean, sem, for 59 looming stimuli; N = 16 animals). D) Example recording of dendritic membrane potential within the LGMD while the eye was exposed to uniform illumination (V_m_ low-pass filtered at 10 kHz and digitized at 40 kHz). E) Despite the high frequency of spontaneous inputs, the V_m_ power is concentrated at frequencies below 5 Hz (black, abscissa is on logarithmic scale). During looming stimuli, the signal power increased 125-fold but remained concentrated at low frequencies (blue). Inset shows the data on a double logarithmic scale. Solid lines and shaded regions are mean ± sem (N = 16 animals). F) Using wavelet analysis to measure the membrane potential power density in both the frequency and time domains reveals that during spiking there is increased high frequency power (dashed square), but the peak power remains at frequencies <10 Hz. Plot displays the average power density map of the same 59 looming responses shown in C and E.

Next, we computed the spectral power density in response to looming stimuli. During the course of such a stimulus the number of activated ommatidia increases from ~10 to over 2,500, producing an increase in activated synapses from ~100 to over 20,000 (Rind et al. 2016). As this increased synaptic input occurs in ever tighter time windows (Fig. 2C), it might lead to a shift in power towards higher frequencies. However, low frequency power increased during looming stimuli, with over 50% of it below 1.5 Hz (Fig. 2E, blue lines). As most of the membrane depolarization and spiking activity generated by looming occurs over the last second before collision, we also used wavelet analysis to resolve the frequency content of the membrane potential in the time domain (see Methods). During this period there was an increase in power spectral density at higher frequencies (Fig. 2F, gray box), but most power remained centered around 1-2 Hz (Fig. 2F). The concentration of the LGMD membrane potential power at low frequencies despite high frequency synaptic input could be due to the activation characteristics of these inputs or to low-pass filtering by the LGMD’s membrane. We tested the latter possibility by measuring the frequency-resolved input and transfer impedance of the LGMD’s membrane. The input impedance reveals the mapping of current to local dendritic membrane potential frequencies, while the transfer impedance documents the change in membrane potential frequency content as the signal propagates from the dendrites toward the SIZ.

### Single and dual dendritic LGMD recordings *in vivo* in response to chirp currents

The LGMD’s input and transfer impedance were measured by recording *in vivo* the membrane potential with sharp electrodes (Fig. 3A). These experiments consisted of injecting chirp currents (sine waveforms with time-varying frequency) during single or paired recordings in the excitatory dendritic field (see Methods). Superposing chirp currents on different holding currents allowed us to measure impedance at multiple steady-state membrane potentials (V_SS_; Fig. 3B), and thus at different steady-state activation levels of subthreshold conductances present in the LGMD membrane. In general, chirp responses looked consistent across V_SS_ values, tapering slightly with increasing frequency. Examination of different periods of the chirp show that at low frequencies the oscillations in membrane current and potential were synchronous, while at higher frequencies the membrane potential lagged behind the current and the change in potential was smaller (Fig. 3C). We conducted similar experiments in voltage clamp and found similar changes to occur (Fig. 3D).

**Figure 3.**
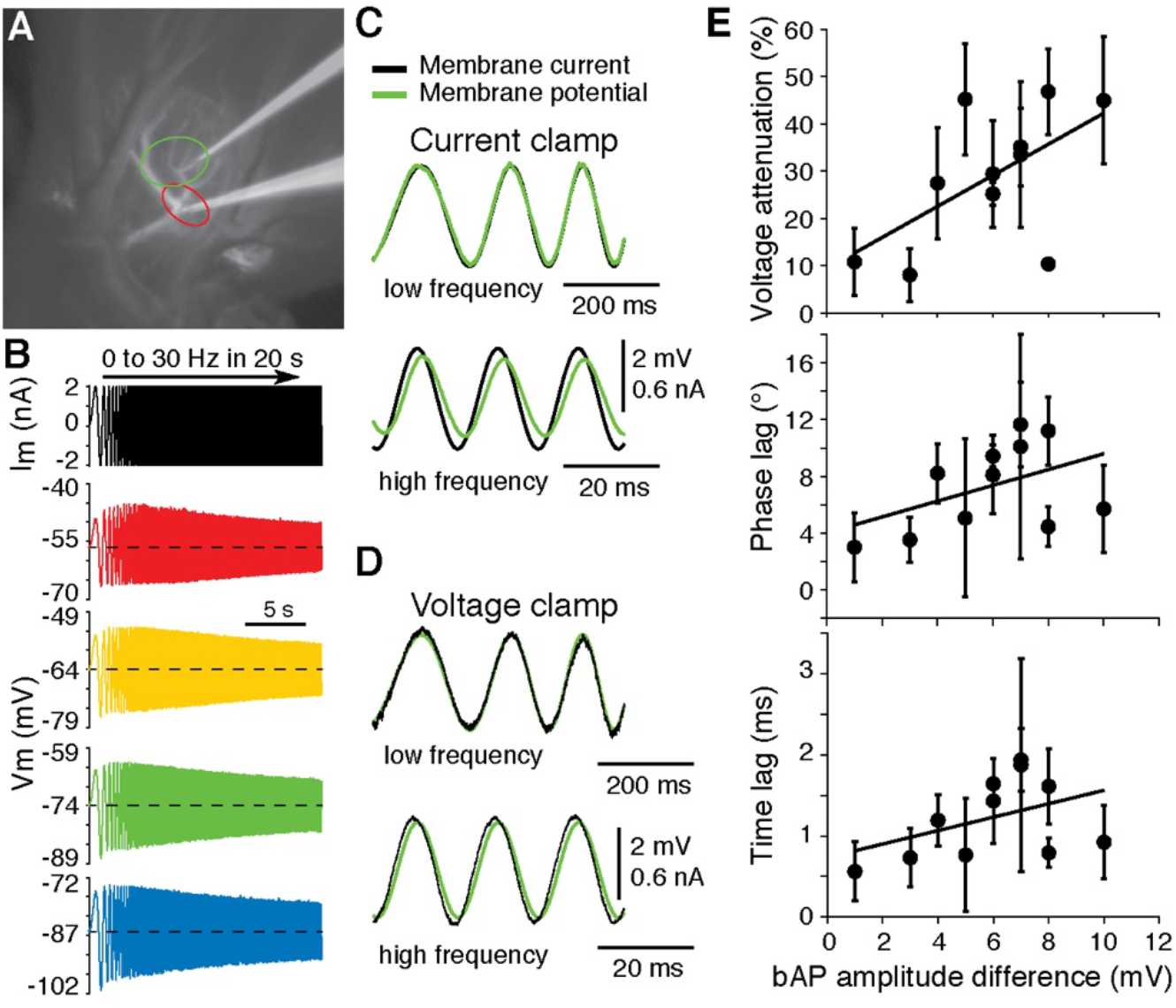
Characterization of LGMD’s membrane impedance profile with *in vivo* dendritic recordings. A) Either single or dual recordings were made from dendritic field A and the primary neurite connecting field A to the SIZ. Image taken with a CCD camera during a dual recording after staining the LGMD with Alexa 594. The internal solution of both electrodes contained Alexa 594 for visualization. Red and green ellipses encompass the region sampled by the proximal and distal electrode, respectively, across 12 recording pairs (8 animals). B) To measure membrane impedance, chirp currents (sine waveforms of increasing frequency) were injected at different steady-state membrane potentials (V_SS_). At top is an example of a linearly increasing chirp current followed by four recordings of the membrane potential in response to this chirp current superposed on different holding currents (I_hold_). The values of I_hold_ and V_SS_ (dashed lines) were 2, 0, −2, −4 nA and −56, −64, −74, −86 mV, respectively (from top to bottom). C) Examples traces of low and high frequency sections of the measured membrane current and potential. At low frequencies the current and potential are synchronous (~0° impedance phase) but at high frequencies the potential trails the current (negative impedance phase). Note that at high frequencies the membrane potential is reduced relative to the current indicating a reduction in impedance amplitude. D) Impedance measurements were similar when measured in voltage clamp. E) Signals attenuated more and had longer lags as the electrotonic distance increased between recording electrodes (estimated by the difference in amplitudes of the backpropagating action potential, bAP). Each point and error bar report the mean value and sd (across different V_SS_ values) for a recording pair (N = 11 recording pairs from 8 animals). Mean bAP amplitudes were 32 and 26 mV for the proximal and distal electrode; the dendritic path distance between recordings ranged from 63 to 149 μm with a mean of 105 μm.

Quantification of transfer impedance characteristics was made as a function of the distance between different pairs of dendritic recordings. The inter-electrode distance was measured both as the dendritic path length between the electrodes in experimental micrographs (path distance, Fig. 3A), and as the difference in back propagating action potential (bAP) amplitudes (electrotonic distance). At the base of the excitatory dendritic field, the bAP amplitude was ~40 mV and decayed to <10 mV in distal dendritic branches. The recording locations in the present study were in larger branches with bAP amplitudes between 20 and 40 mV. The path distances between these electrode pairs ranged from 63 to 150μm. The difference in bAP amplitudes was well correlated with changes in membrane potential transfer properties (Fig. 3E). In contrast, there was no correlation between path distance and voltage attenuation (r(9) = 0.17, p = 0.63, Pearson linear correlation), likely due to differences in dendrite diameter or number of branch points between the recording pairs. With increased electrotonic distance between recording pairs, there was reduced voltage transfer and an increased delay. The closest recordings had ≤10% voltage attenuation, while for more distant pairs this value increased to almost 50% (Fig. 3E). The average phase and time lags between locations were ~3° and 0.5 ms between the most synchronous pairs and ~10° and 2 ms between the most asynchronous pairs. For subsequent analyses of transfer impedance, data from all recording pairs were pooled.

### Consistency of impedance amplitude across holding potentials and chirp frequencies

The impedance amplitude is the absolute value of the complex impedance and represents the gain function, mapping at each frequency a given synaptic or electrode current into a change in membrane potential. We measured the impedance amplitude at different V_SS_ values and examined how its profile changed with holding potential. The LGMD dendrites have a resting membrane potential near −65 mV, can be hyperpolarized by inhibitory inputs to below −70 mV, and are depolarized by excitatory inputs to >-50 mV during looming stimuli (Fig. 2C). We thus used V_SS_ values spanning that range and confined below the spike threshold potential.

Input impedance amplitude was highest at low membrane potentials (<-75 mV), particularly at low frequencies (Fig. 4A, dashed box). Yet, despite the LGMD’s resting HCN and M conductances (Dewell and Gabbiani 2018a, 2018b) changes in V_SS_ produced little change in impedance amplitude (Fig. 4A). This differs from other HCN and M channel-expressing neurons that exhibit a pronounced resonant frequency (Gastrein et al. 2011; Gutfreund et al. 1995a; Hu et al. 2002; Hutcheon et al. 1996). The transfer impedance amplitude similarly changed little with membrane potential and tapered only slightly with increased frequency (Fig. 4B). Accordingly, the voltage attenuation between paired recording locations also showed a high overlap across holding potentials (Fig. 4C). The variability of the voltage attenuation between recording pairs was higher though, due to the differences in electrotonic distance (Fig. 3E). The constancy of impedance amplitude at different V_SS_ values implies a constant total membrane conductance across holding potentials. While other channel types might contribute, HCN and M channels account for the majority of the LGMD’s resting membrane conductance, suggesting that their total conductance remains roughly constant. This in turn would increase the consistency of EPSP amplitudes across holding potentials. This idea was confirmed by injecting currents simulating EPSP waveforms at the same V_SS_ values used to measure impedance, resulting in peak depolarizations that were independent of V_SS_ (r(89) = −0.05, p = 0.63, 91 V_SS_ values from 16 recordings in 9 animals).

**Figure 4.**
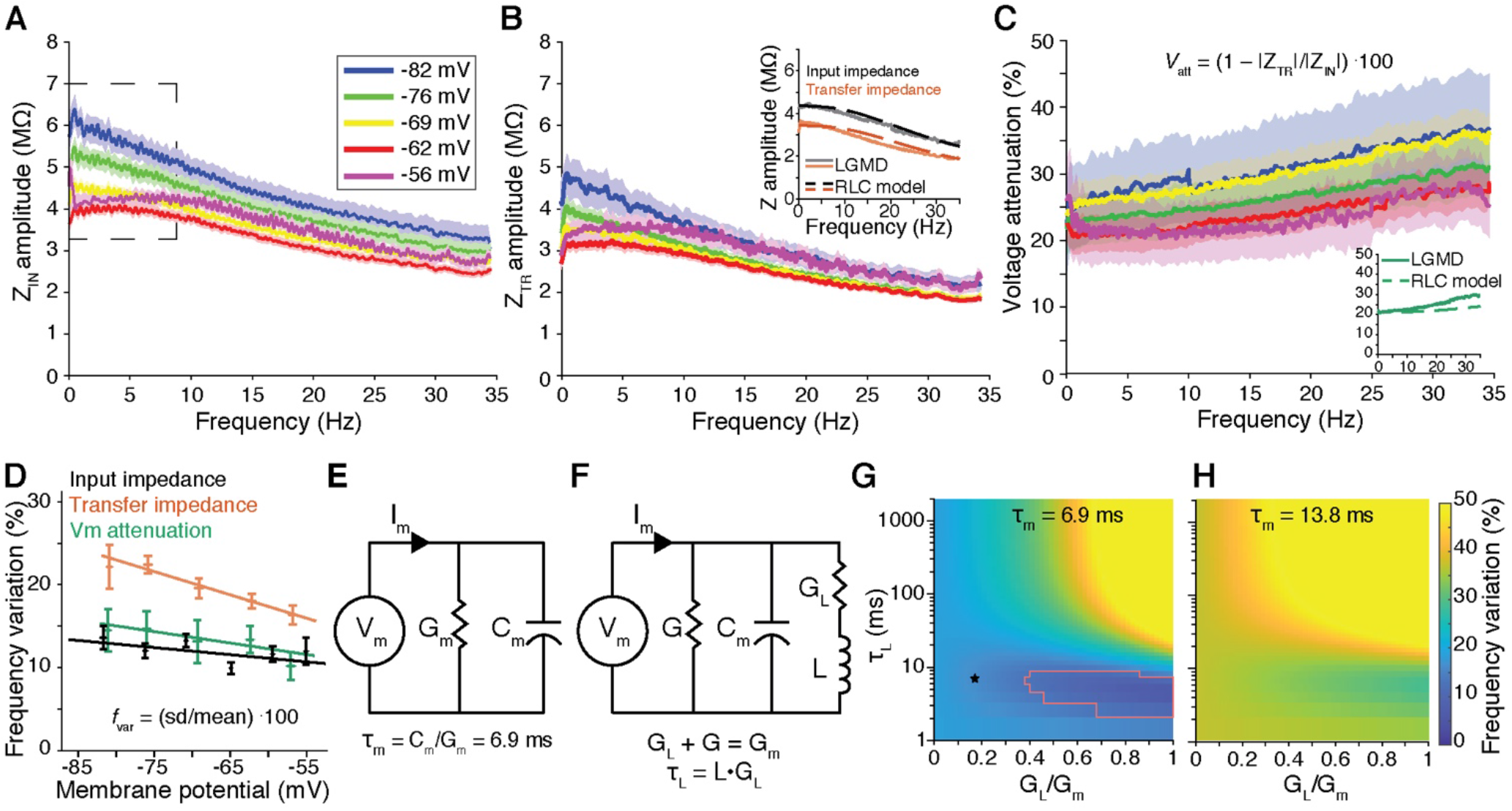
The LGMD membrane impedance amplitude and voltage attenuation behave similarly across frequencies and membrane potentials. A) Input impedance (Z_IN_) amplitude profiles measured at different steady-state membrane potentials (V_SS_, see Fig. 3B) and displayed as mean ± sem (lines and shaded regions) reveal a high degree of overlap across membrane potentials. The lines only show clear separation for low frequencies (<10 Hz, dashed square) at membrane potentials >10 mV below Vrest (65 mV). N = 76 recordings from 56 animals. B) The transfer impedance (Z_TR_) amplitude between different dendritic sites also varied little across membrane potentials. The inset shows the average input and transfer impedance amplitudes (solid lines) overlaid with those of an RLC circuit model as depicted in panel F (dashed lines). N = 13 recording pairs from 10 animals. C) Voltage attenuation (V_att_) is more variable between recordings than impedance amplitude (note the larger sem regions in C when compared to A or B) with ~20–35% reduction in voltage between dendritic locations. The inset compares attenuation with that of the best-fit RLC model of panel F. Same data set as in B. D) The frequency variation (*f_var_*) of the input and transfer impedance amplitude as well as that of voltage attenuation decreased slightly with membrane depolarization. The frequency variation shows the consistency across frequencies, with a value of 0 indicating an ideal resistor. E,F) RC and RLC circuits, respectively, were chained to obtain equivalent cylinder models for comparison with the data. For both circuit models G_m_ = 116 μS/cm^2^ and C_m_ = 0.8 μF/cm^2^. See methods for details on impedance calculations. G) Heatmap illustrating the effect of the RLC circuit inductor parameters on the frequency variation of its input impedance. The left edge (G_L_ = 0) corresponds to the RC circuit depicted in E. Slower inductor time constants (τ_L_) increased frequency variation. Increasing the ratio G_L_/G_m_ increased frequency variability for slow inductors and decreased it for fast inductors. The red line delimits the parameter region that had a frequency variation as low as the LGMD data. The star indicates the parameter values used for the insets in panels B and C which best matched the shape of the LGMD impedance amplitude profiles. H) Heatmap of an RLC circuit with τ_m_ twice that of the LGMD. For this slower circuit model, no inductor parameter yielded an impedance as consistent across frequencies as the LGMD data. In G and H plotted values are for input impedance; transfer impedance maps look similar but with frequency variations of ~1.06 times those in the input impedance maps for all parameters.

The resonance properties of an electrical circuit are characterized by its bandwidth and resonance strength (the Q factor, Horowitz and Hill, 2015). However, neuronal membranes have complex impedance profiles that are not as easily characterized. The most commonly used measure of neuronal resonance strength is the ratio of peak to steady-state (0 Hz) impedance (Koch 1984a). Unfortunately, this measure ignores impedance at other frequencies. So, in addition to this standard measure (see below, Fig. 6H), we wanted an additional measure that incorporated impedance at frequencies below and above the resonant frequency and thus we characterized the variability of the impedance amplitude by its standard deviation across frequencies normalized to the mean impedance amplitude, which we call the frequency variation (see Methods). The frequency variation of the LGMD’s input impedance amplitude was 10-15% and decreased slightly with membrane depolarization (Fig. 4D, black). Transfer impedance amplitude had a frequency variation of ~20% and also decreased with depolarization (Fig. 4D, red). The power spectral density at 0 Hz was 1,000-fold higher than at 35 Hz during looming stimuli and 25-fold higher during spontaneous inputs (Fig. 2E). For the same frequencies, both the input and transfer impedance amplitudes were less than 2-fold higher. For comparison, if the synaptic inputs during looming were broadband and the lowpass spectral density were due to membrane filtering, this would imply a frequency variation >250%. The lack of prominent lowpass filtering rules out the membrane impedance as the cause of the power spectrum concentration at low frequencies (Fig. 2E,F). Instead, this suggests that the low frequency power is due to the activation characteristics of the synaptic input currents during a looming stimulus.

To test what electrical components could produce the impedance amplitude properties of the LGMD, we created an array of RC and RLC circuit models (Fig. 4E, F), as in earlier studies (Koch 1984b; Mauro 1961; Rall 1964). For both model types we used a specific membrane capacitance (C_m_) and conductance (G_m_) yielding a time constant (τ_m_) of 6.9 ms, close to that measured experimentally in the LGMD (Dewell and Gabbiani 2018a, 2018b; Gabbiani and Krapp 2006a; Peron et al. 2007a). The RLC circuit contained, in addition to the parallel conductance and capacitance, a series conductance and inductance to account for the effects of restorative membrane conductances like g_M_ and g_H_ (Fig. 4F). We varied the percentage of G_m_ that was in series with the inductance and the inductance time constant (τ_L_) to determine which, if any, parameters reproduced the LGMD’s impedance amplitude. These two parameters then model the percentage of the membrane conductance passing through the active channels and the time constant of the active conductance. For most of the tested parameter space, the circuit model’s input impedance amplitude varied more with frequency than that of the LGMD (Fig. 4G). Circuits with a fast inductance (≤ 10 ms) had impedance amplitudes as consistent as that of the LGMD, while slow inductances increased the frequency variation beyond that of an RC circuit, which was already higher than that of the LGMD. This is surprising considering that the LGMD’s resting membrane conductance is heavily dependent on rectifying HCN channels with a time constant of ~1 s (Dewell and Gabbiani 2018a). To assess the importance of the LGMD’s membrane time constant to the frequency variation, we examined the same range of inductance values in an RLC circuit with doubled τ_m_ (increasing τ_m_ by increasing C_m_ or by decreasing G_m_ had the same effect). With the higher τ_m_, no inductance parameters produced a frequency variation as low as that of the LGMD. Therefore, the low frequency variation of the LGMD’s impedance profile depends on a fast τ_L_ in addition to the fast τ_m_. Of the LGMD’s known conductances, g_M_ is the most likely candidate to provide such a fast inductance.

### Impedance phase reveals high current – voltage synchrony across membrane potentials

Membrane capacitance produces a phase lag between input currents and their resulting change in membrane potential that increases both with frequency and distance of propagation. Conductances like g_H_ and g_M_ counteract this lag and increase the synchrony between synaptic input currents and the membrane potential (Hu et al. 2009; Narayanan and Johnston 2008; Ulrich 2002; Vaidya and Johnston 2013). To characterize the timing of the LGMD’s membrane potential with respect to current inputs, we thus measured the frequency-dependent dendritic impedance phase. In a passive, isopotential neuron, membrane capacitance forces input impedance phase to saturate at −90° as input current frequency increases (Mauro 1961; Narayanan and Johnston 2008). Conversely, a positive impedance phase indicates that the change in voltage precedes the change in current, a feature requiring a physiological process that resembles an electrical inductance. Like impedance amplitude, the LGMD’s impedance phase varied little with V_SS_ values, distinguishing it again from other neurons (Gutfreund et al. 1995a; Narayanan and Johnston 2008). For all holding potentials, the phase was positive at low frequencies (≤ 1 Hz). Although phase decreased with frequency, it saturated well above −90° and in some recordings even increased at frequencies > 20 Hz. This was the case for both the input and the transfer impedance phase (Fig. 5A and B, respectively). The LGMD subthreshold membrane potential exhibited zero phase and hence was most synchronous with input current at ~1 Hz, in contrast to spiking which is most synchronous with input currents of ~6 Hz (Dewell and Gabbiani 2018b).

**Figure 5.**
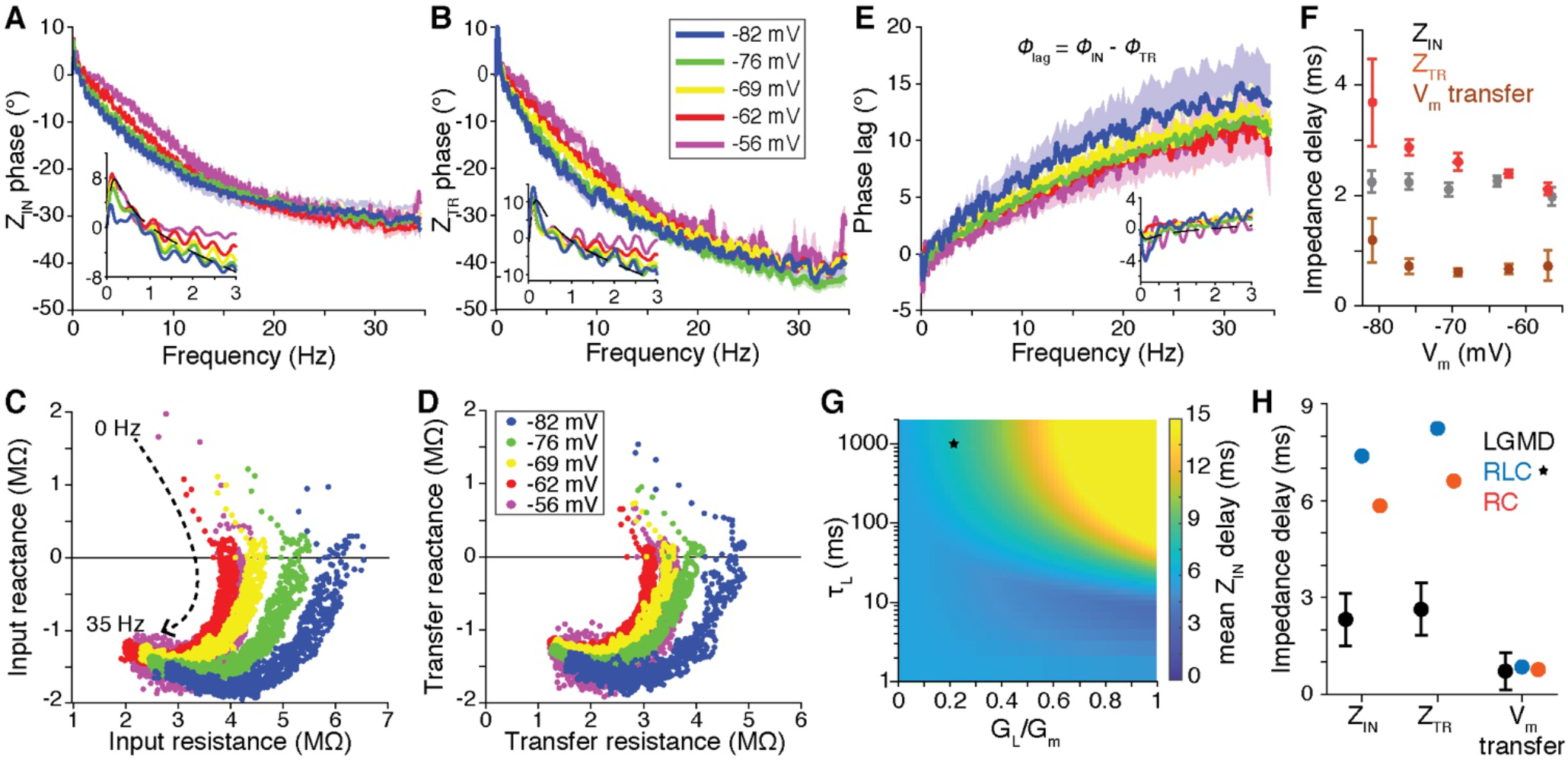
LGMD membrane synchrony and consistency across membrane potentials. A) Z_IN_ phase profiles (mean ± sem) are inductive (exhibit positive phase) only at frequencies < 1 Hz (inset), show high overlap across membrane potentials, and level off around −30°. N = 76 recordings from 56 animals. B) Z_TR_ phases similarly exhibit induction at frequencies < 1 Hz (inset) and overlap across membrane potentials. N = 13 recording pairs from 10 animals. C, D) Impedance locus plots show the real (resistance) and imaginary (reactance) components of the impedance, pooled across frequencies (dotted arrow). The membrane inductance at low frequencies is evidenced by the points with positive reactance. At hyperpolarized potentials, resistance increased but otherwise the LGMD maintained a consistent profile across membrane potentials. E) Phase lag of membrane potential at the ‘recording-only’ location relative to that at the ‘recording and current injection’ location. At frequencies <1 Hz there was a phase advance with the more distant location preceding the input location (inset). The phase lag increased steadily with frequency. F) Impedance delay was measured as the mean absolute time lag between pairs of simultaneously recorded variables. Z_IN_ delay corresponds to the lag between the input current and local membrane potential. Z_TR_ delay was calculated from the lag between the input current and membrane potential at the secondary recording location and V_m_ transfer delay from the lag between the membrane potentials at the two recording locations. Z_TR_ delay decreased with membrane depolarization. G) RLC circuit Z_IN_ delay as a function of its inductor parameters. Z_TR_ delay was ~1.1 times higher than Z_IN_ values shown. The LGMD low frequency inductive phase could only be reproduced by a slow inductor. Dashed black lines in insets of A, B and E produced by parameters marked by black star in G. H) The LGMD exhibited lower delay between membrane current and voltage (phase nearer 0°) than an RC circuit model (G_L_ = 0) or the RLC circuit model marked by the black star in G. No RLC circuit with comparable time constant produced as small a mean Z_IN_ or Z_TR_ delay as the LGMD.

Plotting as in Fig. 5C the real and imaginary components of the input impedance, called respectively resistance and reactance, illustrates both the low frequency inductive and the high frequency capacitive properties of the input impedance phase (corresponding to positive and negative reactance, respectively). Such impedance locus plots combine the information on the impedance amplitude (Fig. 4A) and phase (Fig. 5A): for each point in a locus plot if one were to draw a line segment from the origin to the point, the length of the segment would be equal to the impedance amplitude and the angle relative to the x-axis is the impedance phase. Fig. 5C confirms that the input reactance was consistent across V_SS_ values while the input resistance increased at hyperpolarized potentials resulting in a rightward shift of the curves as V_SS_ decreased. The impedance locus plot of the transfer impedance reveals even less change across membrane potentials, with only a small increase in transfer resistance at hyperpolarized potentials (Fig. 5D). The relative phase of the input and transfer impedances is the membrane potential phase lag with larger values indicating increased transfer delay (Fig. 5E). The phase lag between input and transfer phase was also consistent across V_SS_ values and was negative at frequencies ≤ 1 Hz, indicating an inductive phase shift on the propagating membrane potential. The low frequency inductive characteristics of both the input and transfer impedance reveal the influence of a slow inductor, consistent with the presence of a slowly rectifying HCN conductance.

Signal frequencies with zero reactance (and therefore an impedance phase of zero) have no delay between membrane current and potential. On average, the LGMD’s membrane potential and current were offset by ~2 ms at the input location independent of V_SS_ (Fig. 5F, grey). The mean delay of the transfer impedance was ~3 ms at V_SS_ near −80 mV and decreased with depolarization (Fig. 5F, red). We similarly measured the delay between membrane potentials recorded at two positions, which was ≤ 1 ms (Fig. 5F, brown). To test how this measured timing compared to that of an RLC circuit, we surveyed the same parameter range as depicted in Figure 4. No inductance parameters generated a circuit as synchronous as the LGMD. Circuits with a large and fast inductance generated the lowest input impedance delay, but this was still > 3.5 ms (Fig. 5G). Reproducing the LGMD’s low frequency inductive phase (Fig. 5A,B) required an RLC circuit model with a slow inductance (τ_L_ b≈ 1 s). The dashed black lines of the low frequency insets (Fig. 5A,B & D) were produced by inductance parameters marked by the black star in Fig. 5G. Although the addition of a slow inductance to an RLC circuit reproduced the low frequency inductance of the LGMD membrane, it increased the input and transfer asynchrony of the circuit model further from that of an RC circuit (Fig. 5H). In contrast, the asynchrony between the membrane potential recorded at the two locations could be reproduced by either cable model (Fig. 5H, right). This shows that the LGMD must have an additional synchronizing influence unaccounted for by the circuit model. Whether this increased synchrony is explainable by the LGMD’s active membrane conductances is not immediately evident.

### HCN and M channels reduce impedance amplitude and frequency variation

To assess the influence of HCN and M channels on the LGMD’s input impedance profile, we measured the input impedance before and after addition of two channel specific blockers: ZD7288 and XE991, respectively. Blocking g_H_ reduces the resting membrane potential and input resistance of the LGMD (Dewell and Gabbiani 2018a), so after HCN blockade we applied chirps with lower peak current and higher holding current to generate equivalent V_SS_ values and changes in membrane potential (Fig. 6A). After g_H_ blockade a decrease in the membrane potential was seen with increasing chirp frequencies. The impedance amplitude doubled at low frequency and decreased more steeply with input frequency (Fig. 6B). A reduction in inductance was also clear from the decreased impedance phase (Fig. 6C).

**Figure 6.**
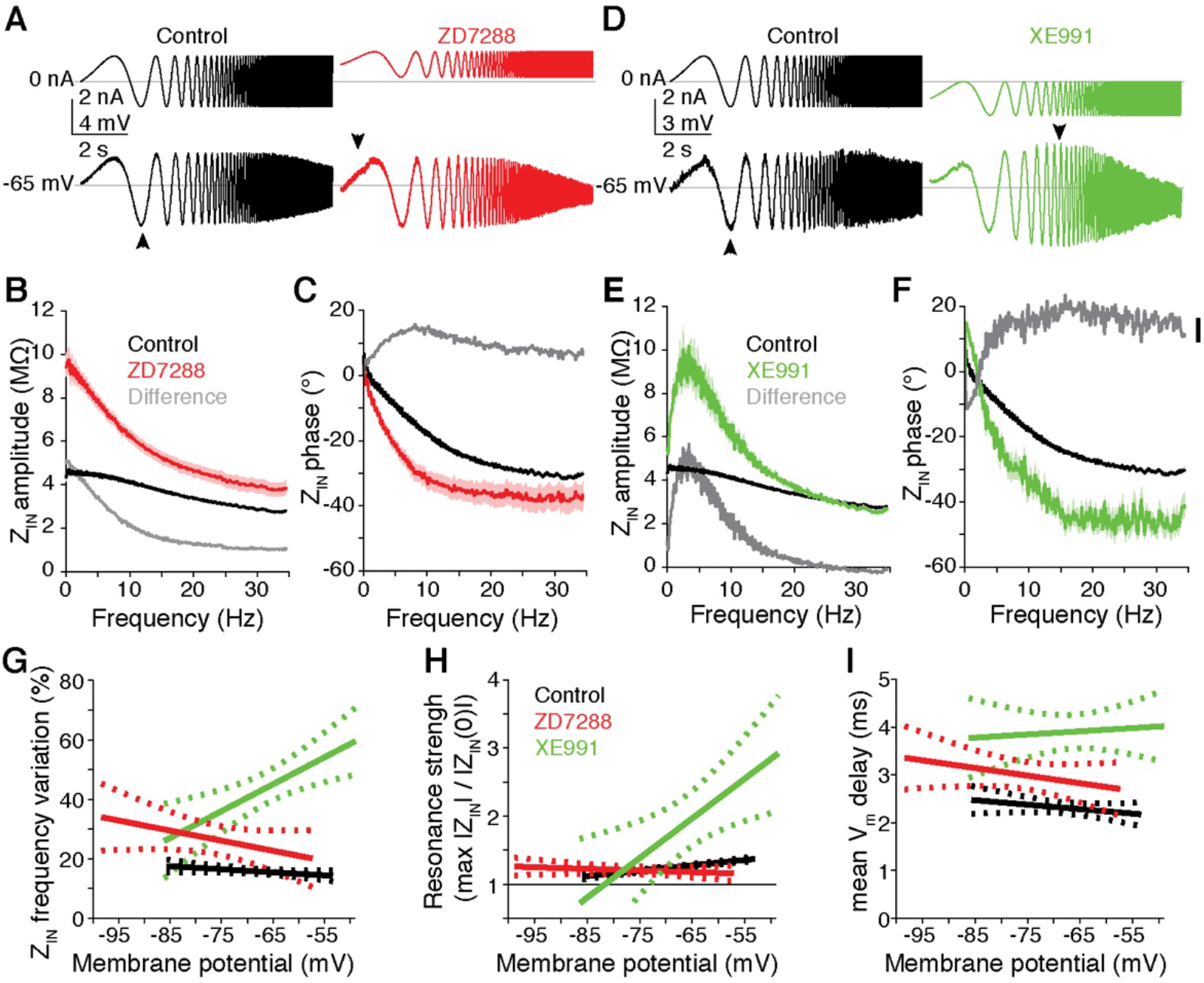
Blockade of either HCN or M channels increased impedance frequency variation and delay between I_m_ and V_m_. A) Example chirp currents (top) and membrane potential responses (bottom) before (left, black) and after (right, red) ZD7288 application. After drug application the chirp amplitude was decreased and I_hold_ was increased to produce a similar V_SS_ and similar changes in membrane potential. B) After blocking HCN channels with ZD7288, input impedance increased 2-fold at low frequencies. Population average impedance (across animals and V_SS_) is shown as mean ± sem (solid line and shaded region). The grey line is the difference between the averages. C) Impedance phase decreased at all frequencies after HCN blockade, reducing membrane current-voltage synchrony. D) Example chirp currents (top) and response (bottom) before (left) and after (right) XE991 application. After drug application the chirp amplitude and I_hold_ were decreased to produce a similar V_SS_ and similar changes in membrane potential. E) After blocking M channels with XE991 a large increase in low frequency input impedance was also seen with peak at ~4 Hz. F) After M channel block impedance phase increased at low frequencies (<2 Hz) but decreased at higher frequencies. G) The increased low frequency impedance after blockade of HCN or M channels produced higher Z_IN_ frequency variations across steady-state membrane potentials. Z_IN_ frequency variation is displayed as linear fit (solid lines) and 95% confidence intervals (dotted lines). M channel blockade increased variation mainly at depolarized potentials while HCN blockade had a greater effect at hyperpolarized potentials. H) The resonance strength, calculated as the maximum impedance amplitude divided by steady-state impedance amplitude (|Z_IN_(0)|), was <1.2 for all membrane potentials under control conditions and after HCN block. After I_M_ block, however, a larger resonance was observed at depolarized membrane potentials. I) The mean absolute time lag between the input current and local membrane potential, increased by ~1 ms after HCN block (red) and ~2 ms after M channel block (green). Control data (black) is the same as shown in Fig. 5F. Control data from 53 recordings at 181 V_SS_ from 44 animals; ZD7288 data includes 9 recordings at 36 V_SS_ from 6 animals; XE991 data includes 6 recordings at 14 V_SS_ from 6 animals.

Blockade of g_M_ increases the LGMD’s resting membrane potential and input resistance (Dewell and Gabbiani 2018b), so we used decreased holding currents and chirp amplitudes after blockade to account for these changes. After g_M_ blockade, a resonance emerged (Fig. 6D, arrowhead). This was reflected in the peak near 4 Hz in the impedance amplitude profile (Fig. 6E). This resonance likely helps generate the increased spiking observed in response to ~4 Hz current inputs following g_M_ blockade (Dewell and Gabbiani 2018b). The impedance phase increased at frequencies ≤ 2 Hz and decreased at higher frequencies (Fig. 6F). These effects suggest that in addition to implementing an inductance evident above 2 Hz, g_M_ also counteracts an additional inductance that increases impedance amplitude at frequencies around 4 Hz and phase at frequencies ≤ 2 Hz.

Both g_M_ and g_H_ reduced frequency variation, causing a larger reduction at depolarized and hyperpolarized potentials, respectively, consistent with their activation kinetics (Fig. 6G). The increased frequency variation after block of either channel was due to a large increase in the range of resistance across frequencies and a reduced high frequency reactance. We also calculated the resonance strength, a common measure of how bandpass a membrane is, and found it to be small at all V_SS_ values in control conditions and after g_H_ block (Fig. 6H, black and red). After g_M_ block, though, at resting V_m_ and above a strong membrane resonance was observed with impedance amplitude around 4 Hz double that of steady state (0 Hz; Fig. 6H, green). Both conductances contribute to decrease the membrane impedance amplitude of the LGMD, with g_H_ causing the largest reduction at and below resting V_m_ and g_M_ at depolarized potentials, respectively. Although we did not measure the effects of g_H_ and g_M_ blockade on transfer impedance, input and transfer resonance were similar in control conditions (with resonant strengths differing by < 0.05 for >75% of trials). So, the changes in transfer resonance after channel block would most likely parallel those of input resonance.

In addition to reducing the impedance amplitude and making it more consistent across frequencies, both g_H_ and g_M_ decreased the delay between input current and the resulting change in membrane potential. After blockade of either channel the input delay increased, with g_M_ blockade having the largest effect (Fig. 6I). After g_M_ blockade the minimum phase was more capacitive at high frequencies and the maximum phase more inductive at low frequencies (Fig. 6F). After g_H_ blockade, the input impedance phase decayed to a minimum at a lower frequency (Fig. 6C). The decrease in input impedance delay produced by g_H_ and g_M_ would improve the LGMD’s ability to discriminate the timing of its synaptic inputs (see below), but even after blocking the channels, the delay between I_m_ and V_m_ was still smaller than any RLC circuit model (compare Fig. 6I and Fig. 5H). This suggests that there must be an additional mechanism beyond these membrane conductances that further reduces these delays, as confirmed by investigating the impact of neuronal morphology (see below).

### Pronounced effect of g_M_ on temporal input discrimination

We have previously shown that blockade of g_H_ reduced the LGMD’s ability to discriminate the spatial pattern of its synaptic inputs (Dewell and Gabbiani 2018a), but the role of the LGMD’s active conductances in discriminating temporal patterns has not been addressed. We tested this with the same sort of simulations described in Figure 1, using a full LGMD model that reproduces most of its experimental properties (Dewell and Gabbiani 2018a, 2018b). Increased noise in the timing of synaptic inputs quickly reduced spiking (Fig. 7A, black). Simulations conducted without g_H_ slightly reduced the model’s sensitivity to input timing, but g_M_’s removal created a substantial reduction (Fig. 7A). The removal of both conductances decreased the ability to discriminate between input timings by over half (Fig. 7A, blue). These changes in spike probability were accompanied by similar effects on spike timing. Removal of g_M_ increased the spike latency (measured from the start of synaptic excitation) as well as its variability (Fig. 7B). This confirms previous experimental findings on the importance of g_M_ in spike timing precision (Dewell and Gabbiani 2018b). In the LGMD, g_H_ plays a critical role in the discrimination of synaptic input patterns during prolonged excitation (Dewell and Gabbiani 2018a), but the present simulations indicate that it does not play a similar role for brief periods of synaptic excitation.

**Figure 7.**
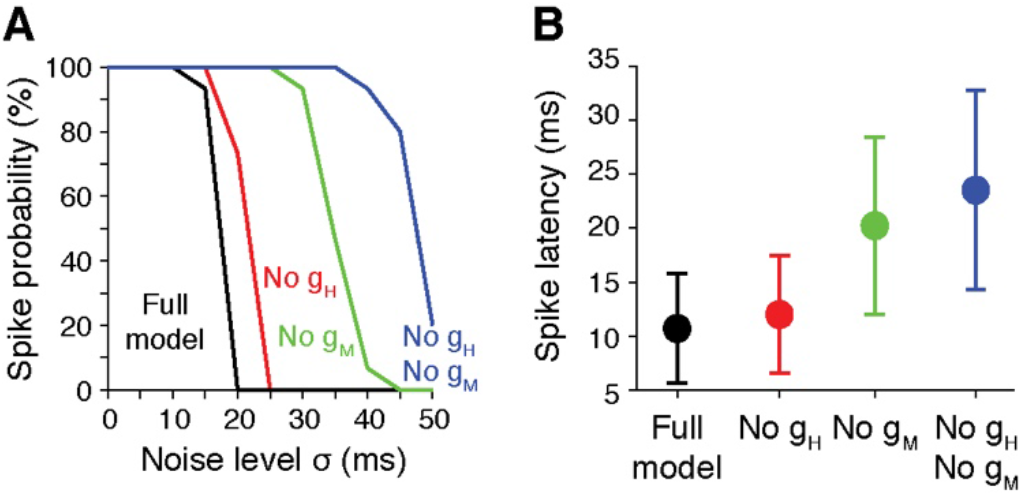
LGMD model simulations show g_H_ and g_M_ increase sensitivity to synaptic input timing and reduce spike latency. A) In the full LGMD model (black), synchronous inputs reliably generated spikes, but for inputs with less reliable input timing (σ > 15 ms) no spikes were produced. Removal of g_H_ (red) slightly decreased the input timing selectivity, and removal of g_M_ (green) or both conductances (blue) produced larger reductions in the timing discrimination. In each condition the leak reversal potential was adjusted to maintain a constant resting membrane potential. B) The timing of the spikes generated were also less reliable. The full LGMD model generated spikes with the shortest latency after the inputs began. After g_M_ blockade spikes occurred later and with less reliable timing.

### Neuronal morphology increases membrane gain and high frequency synchrony

To test the influence of morphology on membrane impedance we compared simulations between passive models that were either isopotential or had realistic branching and attenuation. These simulations build upon earlier ones that examined the role of morphology on signal propagation and impedance in other neurons (Holmes et al. 1992; Jaffe and Carnevale 1999; Mainen and Sejnowski 1996; Vetter et al. 2001). The principal difference in the current simulations is the characterization of morphological influence on the impedance delay and calculation of the net influence of morphology on input currents distributed throughout the dendritic arbors.

Plotting the input impedance delay of every section of the full LGMD model (Fig. 8A) revealed that it was reduced from that of an isopotential model (dashed line) indicating higher synchrony between a local input current and membrane potential. For the same frequency range used in experiments, the passive model segments had a mean input delay of 2.1 ± 1.1 ms, matching experimental data (c.f. Fig. 5H). This suggests that the LGMD morphology plays a bigger role in the membrane potential’s delay relative to input current than its active conductances. The input impedance amplitude was also altered by cell morphology, decreasing less with frequency than in an isopotential equivalent model (Fig. 8B; solid vs. dashed lines). The input frequency variation of the model sections was 8.6 ± 4.7% for the physiologically dominant frequency range (< 35 Hz; see Fig. 1), demonstrating that the LGMD’s morphology alters the input impedance amplitude profile as much as its active conductances. At higher frequencies, the input impedance amplitude remained monotonically decreasing, but was always higher than that of the isopotential model (Fig. 8B, dashed line).

**Figure 8.**
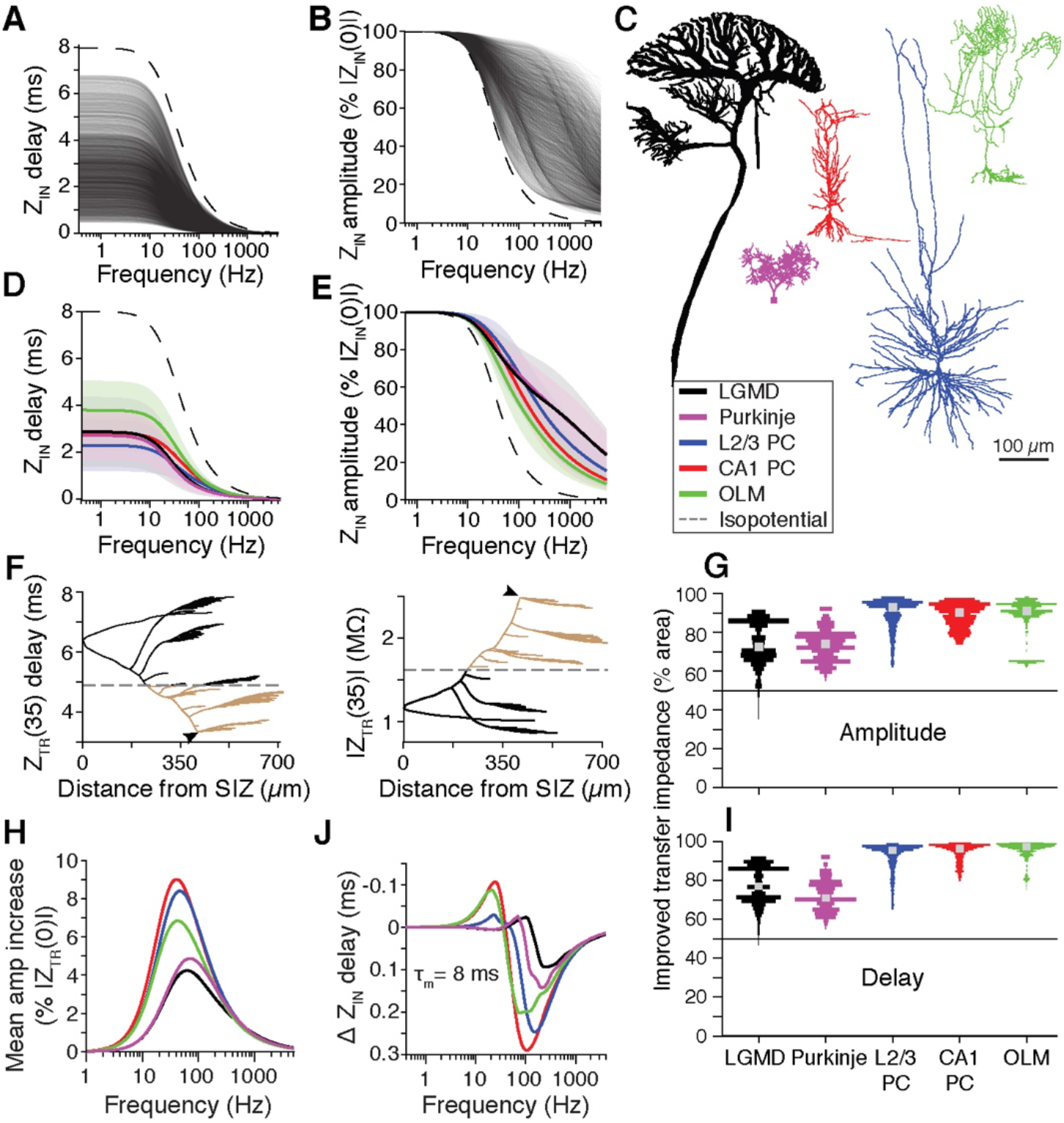
Simulations of passive neurons reveal the influence of morphology on impedance characteristics. A) Every section of the LGMD model (thin black lines) had shorter input impedance delays than an isopotential model (dashed line). B) All sections of the LGMD model (thin black lines) exhibited a decrease in input impedance amplitude smaller than that of an isopotential model (dashed line). For each section the impedance amplitude is normalized to impedance at 0 Hz (|Z_IN_(0)|). The difference between the input impedance amplitude and that of an isopotential model was maximal at ~200 Hz. C) The effect of neuronal morphology on impedance was tested in the LGMD (black) and 4 other cell morphologies of different size and shape: a cerebellar Purkinje cell (purple), a hippocampal CA1 pyramidal cell (red), a cortical layer 2/3 pyramidal cell (blue), and a CA1 oriens lacunosum-moleculare interneuron (OLM; green). D) All neuron models tested had large decreases in input delay compared to an isopotential equivalent model. Data is color coded as depictions in C and presented as mean ± sd (solid lines and shaded regions). E) All morphologies had more consistent input impedance amplitude relative to their isopotential equivalents, with lower frequency variation. Data presented as in D. F) Data from example dendritic segments of the LGMD model shows that at 35 Hz the transfer impedance delay (left) increased and the amplitude (right) decreased with distance from the site of current injection (arrowheads). Segments for which transfer impedance had higher amplitude or shorter delay than that of an isopotential equivalent neuron (dashed line) are brown. Data in G and I measured by summing the surface area of brown segments and dividing by the total surface area. G) For each neuron model segment, we summed the surface area of other model segments for which signal transfer was increased compared to an isopotential equivalent and divided by the neuron’s total surface area. Bar widths indicate how many model segments had improved transfer (higher gain and lower delay) for each percentage range, and grey squares mark the average segment. All morphologies had increased transfer amplitudes compared to an isopotential equivalent model, with the LGMD and Purkinje morphologies having improved transfer for a smaller percentage of the neuron segments. H) For all models, an average all-to-all transfer impedance was measured as a function of frequency and compared to the impedance of an isopotential equivalent. The increase of each cell’s mean transfer amplitude as a function of frequency from isopotential reveals a bandpass increase in average transfer gain. I) The percentage of membrane area with reduced transfer delay was similar for each morphology as that with an increase in transfer gain. Plotted as in G. K) The average all-to-all transfer delay from I_m_ to V_m_ for each model was similar with no difference at low frequencies, a slightly longer delay near the frequency of the cell’s peak gain, and reduced delays at higher frequencies.

To determine whether the effects of morphology on impedance was unique to the LGMD, we conducted similar simulations using four other neuronal morphologies covering a range of sizes and branching patterns (Fig. 8C). The additional morphologies examined were taken from cerebellar Purkinje, layer 2/3 pyramidal, CA1 pyramidal, and oriens-lacunosum moleculare (OLM) neurons. The membrane conductance and capacitance were uniform and the same for all models producing a membrane time constant of 8 ms. Simulations with slower τ_m_ yielded qualitatively similar results. For all morphologies, input impedance delay decreased (Fig. 8D) and amplitude increased (Fig. 8E) relative to their isopotential equivalents (otherwise identical models with zero axial resistance).

Hence, the branched structures of the LGMD and other neurons enhance local inputs with increased gain and higher synchrony between input currents and changes in membrane potential. This is offset in part by lower signal transfer because the branching and axial resistance also increase voltage attenuation between cell regions. The net effect of the morphology on the membrane impedance is therefore not immediately apparent. In experiments, transfer impedance was measured by injecting current at a single location and recording the membrane potential at another. Synaptic currents, however, impinge at thousands of locations spread out across the dendrites. Furthermore, these inputs don’t propagate independently to the site of spike initiation.

The precise effect of morphology on synaptic integration depends on the pattern of its inputs, which are unknown for most neurons, so we assumed broadband inputs spread across the entire arbor. To assess the net effect of morphology on impedance we thus calculated the transfer impedance in an all-to-all manner, with current frequencies between 0 and 5 kHz injected successively into each model segment and the impedance measured in all other segments of the model. An example of transfer impedance profiles from one dendritic segment of the LGMD is illustrated for a 35 Hz current in Fig. 8F (delay on left, amplitude on right). These plots show that segments near the current injection site (arrowheads) had higher transfer impedance while more distant segments had lower transfer impedance than an isopotential equivalent (dashed lines). For all segments of all models, at all frequencies, nearby segments exhibited increased and distant segments decreased transfer impedance relative to the isopotential models. To determine how far signals propagated before the increased attenuation overcame the increased input impedance, we quantified, for each segment, its area of increased transfer impedance relative to an isopotential equivalent. This was recorded as the percentage of membrane for which transfer amplitudes were higher and delays shorter compared to the equivalent isopotential model. In Fig. 8F, this percentage would correspond to the surface area of the segments marked in brown divided by the total membrane area (brown and black).

For currents injected into some LGMD segments, the surface area with increased transfer amplitude was as high as 90% and amounted to ~70% for the average segment (Fig. 8G, black). The Purkinje model had a similar area of increased transfer as the LGMD, while models of the pyramidal and OLM neurons, however, had an increased transfer amplitude from most segments to ~90% of the neuron’s surface area (Fig. 8G). In addition to quantifying the area of improved transfer, we also calculated membrane area weighted averages of the transfer amplitude to examine the net effect of the morphology on signal transfer. Examination of the net transfer amplitude at different frequencies showed that each morphology generated a bandpass increase in transfer amplitude (Fig. 8H). The CA1 pyramidal model had the highest increase, with 40 Hz inputs increasing by 9%. The LGMD and Purkinje morphologies generated smaller increases, 4-5%, shifted at higher frequencies near 65 Hz. This shows that the branching morphologies of neurons provide a bandpass increase to the overall membrane gain. As branched morphologies cause membrane potentials to attenuate, it would not have been surprising if their net effect were a reduced transfer amplitude relative to an isopotential model, but this was not the case.

For all morphologies, the reduction in input delay was partially offset by the increase in transfer delay. The improved timing of the input impedance spread throughout most of the neurons, and the lower transfer delay propagated to almost the same membrane area as did the increased transfer amplitude (Fig. 8I). To calculate the total transfer delay, values were weighted by the strength of the transfer amplitude: if there is minimal signal transfer between 2 segments then the timing of that transfer is less relevant. The net impact of morphology on the average transfer delay was small (≤ 0.3 ms for all frequencies; Fig. 8J). All morphologies had slightly smaller net transfer delays than an isopotential equivalent model for signals above 100 Hz. The amount of this improvement scaled with the τ_m_ used for simulations; simulations with τ_m_ = 16 ms had twice the improvement in delay due to morphology. The overall effect of the morphologies on transfer timing was minimal on low frequency signals, but could improve the timing of high frequency signals, like those generated from fast synaptic currents.

## Discussion

Here we have described the impedance properties of the LGMD *in vivo*, measured over the range of frequencies encompassing most of the signal power mediated by the LGMD’s synaptic inputs. We found that the membrane impedance amplitude of the LGMD is consistent over the range of subthreshold membrane potentials and input frequencies involved in the detection of approaching objects. Further, the membrane impedance revealed small delays between input current and resulting changes in membrane potential. This membrane timing and gain consistency were both shaped by the conductances g_H_ and g_M_, as well as the neuron’s branching morphology. Extensive modeling revealed that these changes to the LGMD’s impedance profile increase its ability to discriminate the temporal patterns of excitatory synaptic inputs.

Despite a high rate of spontaneous and stimulus-evoked synaptic inputs due to the large number of excitatory synapses the LGMD receives (Rind et al. 2016), most of its membrane power was concentrated at low frequencies (Fig. 2E). During looming stimuli, as the rate of excitation increased, the dominant signal frequency actually decreased with over 50% of the membrane potential spectral power below 1.5 Hz (Fig. 2E, F). As neither the input nor the transfer impedance amplitude of the LGMD were lowpass over the same frequency range (Fig. 4A, B), this must reflect the characteristics of pre-synaptic inputs to the LGMD. Neither the membrane gain, nor the delay between I_m_ and V_m_ changed dramatically over the frequencies tested, favoring integration over a broad frequency range during looming.

To determine what shapes the frequency dependence of the impedance gain we used pharmacological manipulations and computational modeling. We found that the neuron’s morphology and the active conductances g_H_ and g_M_ (Fig. 6G) dramatically reduced frequency variation of the impedance gain, making the LGMD membrane impedance more broadband. This was surprising, as both g_H_ and g_M_ usually make neurons more bandpass (Hönigsperger et al. 2015; Hu et al. 2009; Hutcheon and Yarom 2000; Narayanan and Johnston 2008). The primary effect of g_H_ on the membrane impedance (Fig. 6B,C), is actually the same as in other neurons, a reduction of lowpass filtering and impedance delay. Whether g_H_ narrows the membrane bandpass properties or broadens them, as it does here and in CA3 interneurons (Anderson et al. 2011), depends on the relative balance of its effects and those of other membrane properties.

In addition to showing low frequency variation and delays between I_m_ and V_m_, the impedance of the LGMD also showed consistency across holding potentials. As an object approaches, increasing excitation causes dendritic depolarization of >20 mV, so changes in dendritic integration properties with membrane potential would influence the detection of impending collisions. Neither the amplitude nor the phase of the membrane impedance changed much from −70 to −55 mV (Fig. 3 and 4). This consistency can also be largely explained by g_H_ and g_M_. At hyperpolarized potentials g_H_ increases, dominating the membrane conductance, while conversely g_M_ increases with depolarization. At the LGMD’s resting membrane potential (−65 mV), both channels have large conductances and shallow activation kinetics (Dewell and Gabbiani 2018a, 2018b). If the activation ranges of g_H_ and g_M_ had less overlap, the impedance profile would be less consistent across membrane potentials and the channels would be more likely to create separate hyperpolarized and depolarized resonances as seen in pyramidal cells (Hu et al. 2002, 2009).

Synaptic integration is influenced by the input timing as well as its gain. We found that the LGMD exhibited shorter delays between I_m_ and V_m_ than could be explained by a circuit model (Fig. 5H). These delays were reduced by both g_H_ and g_M_ (Fig. 6I). On average, blockade of g_M_ produced a larger reduction than g_H_, except at frequencies near the peak membrane power during looming stimuli (~1.3 Hz; Fig. 2F). The influence of g_H_ at low frequencies is consistent with its effect on synaptic pattern discrimination during looming (Dewell and Gabbiani 2018a), while having only a small effect in model simulations with brief periods of excitatory inputs (Fig. 7A). In contrast, the ability to discriminate the timing of these simulated synaptic inputs was greatly reduced by removal of g_M_ from the LGMD model (Fig. 7), a feature that would be difficult to test experimentally because the g_M_ blocker XE991 also has presynaptic effects (Dewell and Gabbiani 2018b). Previously, simulations of these same brief periods of excitation using a model of a CA1 pyramidal neuron found g_H_ to dramatically increase sensitivity to input timing (Migliore et al. 2004). In those CA1 simulations the g_H_ time constant was ~40-fold faster than in our LGMD model, but it was very similar to the LGMD’s g_M_ time constant. This further supports the idea that these effects are mainly due to the change in membrane impedance, rather than channel type.

Although g_H_ and g_M_ decreased I_m_-V_m_ delays, the delays remained lower after their blockade than those of an equivalent circuit model. Testing a morphologically realistic, passive LGMD model revealed that the neuron’s shape influenced the membrane’s input impedance as much as the active conductances (Fig. 8A, B). The branching morphology introduces new electrical paths whose axial resistivity does not decrease with frequency. This led to increased input gain and reduced input delays. Previous modeling of pyramidal and Purkinje neurons have also found that dendritic branches reduced lowpass filtering (Dhupia et al. 2015; Ostojic et al. 2015). This led us to wonder how the influence of LGMD’s morphology on impedance compared to other cell morphologies, and whether the increase in input impedance caused by an extended neuronal morphology was counteracted by decreased signal propagation. We tested this with transfer impedance simulations of five neuronal morphologies.

Transfer impedance is defined between two locations. While it is technically infeasible to simultaneously record intracellularly from many places within a neuron, in a model it is no more difficult to measure transfer between one hundred locations than between two. The reason we chose to measure the all-to-all transfer rather than just transfer from individual dendrites to the site of spike initiation is simple. In neurons, *in vivo* synaptic currents impinge throughout the dendrites and do not propagate in isolation, but continuously integrate with new inputs. As far as we know, this is the first examination of the sum transfer of a dendritic arbor in this fashion. It revealed that in the LGMD, as well as other cells, the net effect of morphology is an increase in all-to-all impedance gain and a decrease in high frequency delay between I_m_ and V_m_. Additionally, we found that the net increase in gain is bandpass between 30 and 100 Hz (Fig. 8H). Although their magnitudes were consistent, the frequency range of these changes depended on the simulated membrane time constant; for simulations with a slower membrane the bandpass gain increase shifted to lower frequencies (data not shown). This furthers previous findings on the effects of dendritic morphology on a neuron’s integration capabilities (Dhupia et al. 2015; Mainen and Sejnowski 1996; Moore et al. 1988).

Ultimately the integration properties of a neuron are dictated by the pattern of its inputs and its computational role. For hippocampal pyramidal neurons that receive theta rhythmic inputs, a strong theta (4-10 Hz) resonance and synchrony could be advantageous (Hu et al. 2009; Hutcheon and Yarom 2000; Vaidya and Johnston 2013). Similarly, it might be a computational benefit for cortical neurons to exhibit large impedance changes with membrane potential to enhance differences between up and down network states (Gutfreund et al. 1995b; Hasselmo 2005; Heys et al. 2010; Wang 2010). For auditory neurons, specialized in detecting high frequency sound waves, a very fast membrane time constant and high frequency inductance lead to broadband membranes with resonance at frequencies up to 400 Hz (Mikiel-Hunter et al. 2016; Remme et al. 2014). The LGMD detects approaching objects and discriminates their specific input statistics from a wide range of other input patterns through their integration over a wide range of membrane potentials. This involves discriminating synaptic input patterns on the order of milliseconds to tens of milliseconds (Jones and Gabbiani 2010) as well as over several seconds (Dewell and Gabbiani 2018a). Thus, achieving broadband membrane impedance with minimal delay across input frequencies and membrane potentials by a balance of active membrane conductances and an extended morphology is exactly what is needed to fine tune escape behaviors.

